# Heme Attenuates Endogenous Opioid Levels in Leukocytes of HIV positive individuals with Chronic Widespread Pain

**DOI:** 10.1101/2020.04.24.059790

**Authors:** Saurabh Aggarwal, Jennifer J DeBerry, Israr Ahmad, Prichard Lynn, Cary Dewitte, Simran Malik, Jessica S Merlin, Burel R Goodin, Sonya L Heath, Sadis Matalon

## Abstract

The prevalence of chronic widespread pain (CWP) in people with HIV (PWH) is high, yet the underlying mechanisms are elusive. Leukocytes synthesize the endogenous opioid, β-endorphin (β-END), within their endoplasmic reticulum (ER). When released into plasma, β-END dampens nociceptive transmission by binding to opioid receptors on sensory neurons. In the present study, we hypothesized that heme-induced ER stress attenuates leukocyte levels/release of β-END, thereby increasing pain sensitivity in PWH. Results demonstrate that PWH with CWP have fragile erythrocytes, high plasma levels of cell-free heme, and impaired heme metabolism. Leukocytes from PWH with CWP also had high ER stress and low β-END compared to PWH without CWP and HIV-negative individuals with or without pain. *In vitro* heme exposure decreased β-END levels/secretion in murine monocytes/macrophages, which was prevented by treatment with sodium 4-phenylbutyrate, an ER stress inhibitor. To mimic hemolytic effects in a preclinical model *in vivo*, C57BL/6 mice were injected with phenylhydrazine hydrochloride (PHZ). PHZ increased cell-free heme and ER stress, decreased leukocyte β-END levels and hindpaw mechanical sensitivity thresholds. Treatment of PHZ-injected mice with the heme scavenger, hemopexin, blocked these effects, suggesting that heme-induced ER stress and a subsequent decrease in leukocyte β-END may contribute to CWP in PWH.

## BACKGROUND

In the modern treatment era, individuals infected with human immunodeficiency virus-1 (HIV-1) who are diagnosed and treated early can have a near-normal life expectancy. However, chronic widespread pain (CWP) in people with HIV-1 (PWH) is associated with a high rate of disability and reduced quality of life (1), with prevalence estimates ranging from 25%-85% (2-4).

Leukocytes (neutrophils, monocytes/macrophages, and lymphocytes) are a rich source of endogenous opioid peptides (5-13) that inhibit nociceptive transmission by binding to peripheral opioid receptors (14-16). During inflammation, leukocytes are recruited to the site of damage (17). Upon stimulation with mediators such as interleukin-1 (IL-1), corticotropin-releasing factor (CRF), or norepinephrine, they release opioid peptides (i.e., β-endorphin (β-END) which exert an anti-hyperalgesic effect in inflamed tissues (12, 18-22), (18, 19, 21, 22). Blocking the action of endogenous opioids with antibodies or elimination of peripheral immune cells that produce and release them has been shown to decrease this effect (14, 18, 22). Preclinical studies support a role for immune cells in endogenous analgesia. For example, drug-induced immunosuppression in rats has been shown to increase mechanical and thermal hyperalgesia during inflammation (23), and adoptive transfer of allogenic polymorphonuclear leukocytes (PMNs) after innate PMN depletion restores opioid receptor-mediated analgesia during inflammation (24). Macrophages, in particular, generate and release opioid peptides in response to inflammation or injury, with greater opioid content and release by the M2-polarized, pro-resolution phenotype than the pro-inflammatory, M1 phenotype (25). Promoting the polarization of naive macrophages toward the M2 phenotype (26, 27) can attenuate clinical postoperative pain (28) and decrease tactile hypersensitivity in animals (29).

Encapsulated heme is a ubiquitous molecule that plays an essential role in various physiological functions. However, when liberated from red blood cells (RBCs), cell-free heme can have harmful consequences. Previous studies by our group and by others have shown that cell-free heme is a pro-inflammatory molecule that causes endoplasmic reticulum (ER) stress and cellular injury (30-33). Heme activates toll-like receptor 4 (TLR4) signaling in macrophages (34, 35), causing the release of pro-inflammatory, pro-algesic cytokines (e.g., IL-1α, IL-6, TNF-α), chemokines, and inducible nitric oxide synthase (iNOS) (36-38). In macrophages, heme-induced ER stress promotes apoptosis and polarization to the pro-inflammatory, M1 phenotype (32, 39, 40). Clinical and preclinical studies have shown that cell-free heme is correlated with acute painful vaso-occlusive crises in children with sickle cell disease and in transgenic sickle mice (41, 42). Animal studies have demonstrated that heme oxygenase-1 (HO-1), a heme metabolizing protein, ameliorates hyperalgesia due to nerve injury (43) and inflammation (44). However, whether cell-free heme is directly involved in chronic pain is not known.

The objective of the present study was to examine the role of cell-free heme as a mediator of CWP in PWH, since HIV infection is associated with susceptibility to hemolysis due to secondary sequelae (45-50). Specifically, we hypothesized that increased cell-free heme impairs peripheral endogenous analgesic mechanisms by increasing ER stress in peripheral immune cells, leading to increased pain. Using a multifaceted approach, the current study provides evidence that cell-free heme contributes to CWP in PWH and characterizes a mechanism through which it exerts this effect.

## RESULTS

### PWH who self-report CWP have increased plasma levels of cell-free heme and impaired heme scavenging

As a first step in establishing cell-free heme as a contributing factor to HIV-associated chronic pain, plasma concentrations of cell-free heme, RBC membrane oxidation and fragility, and the heme scavenging protein, hemopexin (Hx), were measured in HIV-positive and -negative people with or without CWP. The most common sites of pain reported by PWH are low back (86%), hands/feet (81%), and knee (66%) (51); therefore, HIV-negative individuals with chronic low back pain (LBP) were included as a second control group in addition to PWH without chronic pain.

General demographic characteristics of participants were not significantly different among groups (age range 45-52 years; race distribution 66-68% African American), with the exception of sex (M>F) (**Table 1**). Reflective of the patient population of the clinic from which HIV-positive individuals were recruited, the proportion of men in the HIV-positive group was 52% vs. 33% in the HIV-negative groups. De-identified clinical data showed that the average current and nadir CD4^+^ cell counts and average current and highest viral load (VL) were not different between HIV-positive individuals with or without CWP.

**Table 1.**
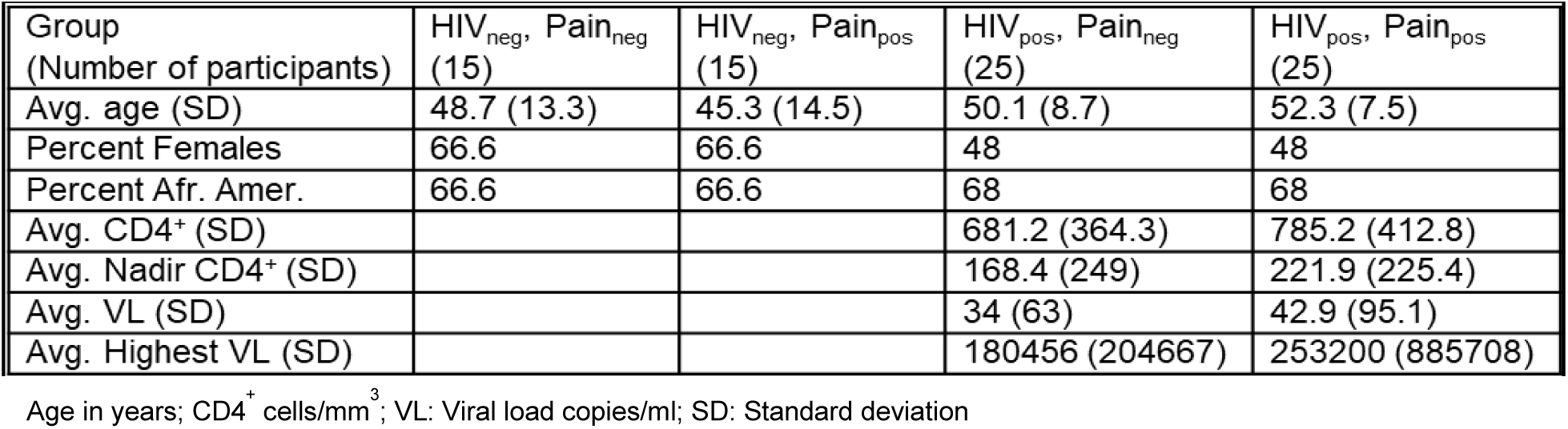
Demographic and clinical data of participants. The table shows data obtained from HIV positive or negative individuals with or without pain: average current and nadir CD4^+^ cell count, and the average current and highest viral load (VL).

Whole blood samples were collected from study participants for analysis of plasma levels of total (free and protein-bound) cell-free heme. Data demonstrated that PWH with CWP had significantly higher (2-3 fold increase) cell-free heme levels than other groups (**Figure 1A**). However, Hx concentrations were comparatively lower in PWH with than without CWP, suggesting that heme scavenging machinery is attenuated in the context of pain (**Figure 1B**). *Ex vivo* RBC mechanical fragility was assayed by quantification of heme/hemoglobin (Hb) release in response to mechanical stress using freshly obtained RBCs from participants (52). Data demonstrated that PWH had significantly higher RBC fragility, resulting in hemolysis, compared to HIV-negative individuals (**Figure 1C**). Further, hemolysis was also greater in PWH with CWP compared to PWH without chronic pain. In addition, RBC membranes were isolated for measurement of carbonyl (aldehyde and ketone) adducts, a hallmark of protein oxidation. Protein carbonylation in RBCs was significantly higher in PWH with CWP compared to the HIV-negative individuals with LBP and both pain-free groups (**Figure 1D**). Together, these results suggested that PWH with CWP were prone to hemolysis and thus exhibited elevated plasma levels of cell-free heme. All PWH (with or without pain) were on anti-retroviral therapy (ART) and had low viral copies/ml of blood) suggesting that RBC hemolysis and elevated cell-free heme are not dependent on VL or ART.

**Figure 1.**
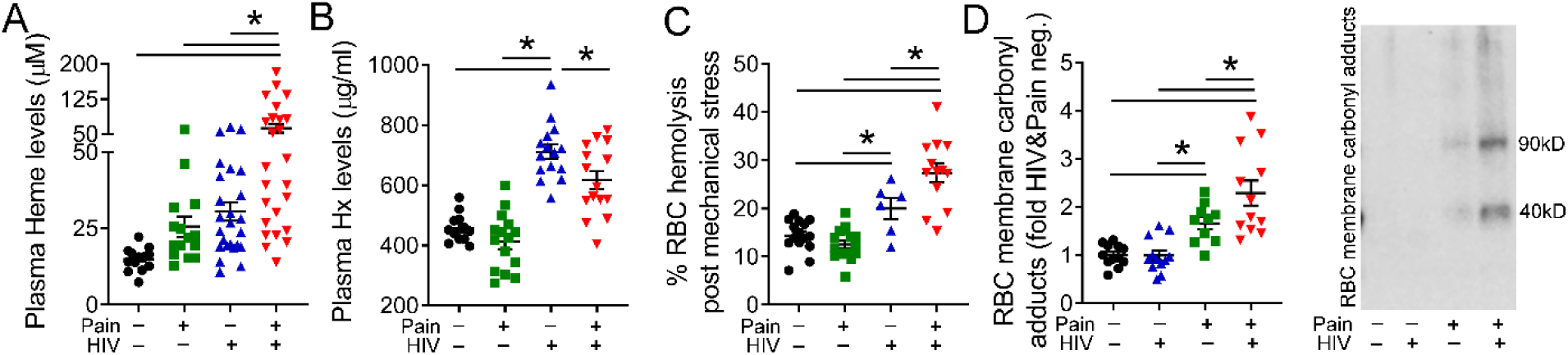
PWH with CWP have increased plasma levels of cell-free heme and impaired heme scavenging. Total (free and protein bound) cell-free heme was measured in plasma. PWH with CWP had significantly higher heme than HIV-1 negative controls and PWH without pain (n=15-25) (A). Plasma hemopexin (Hx) levels were lower in PWH with CWP than PWH without pain (n=12-15) (B). PWH with CWP had increased RBC hemolysis compared to other groups (n=6-25) (C). Protein carbonylation in RBC membrane was significantly higher in PWH with CWP compared to the no-pain groups and the HIV-negative group with LBP (n=10-12) (D). Individual values and means ± SEM. \**P* < 0.05 vs. groups at the end of individual lines; one-way ANOVA followed by Tukey post-hoc testing.

### PWH with CWP exhibit a pro-algesic immune profile

Immune cells release both pro-inflammatory, algesic, and anti-inflammatory, analgesic mediators, the balance of which, contributes to the presence of hyperalgesia or allodynia. We have previously demonstrated that heme increases pro-inflammatory cytokines (33). In the current study, plasma profile of 10 cytokines was quantified in the participants. Data showed that PWH with CWP exhibited significantly higher levels of proalgesic cytokines including IL-1β (**Figure 2A**), IL-6 (**Figure 2B**), and TNF-α (**Figure 2C**), that are produced predominantly by pro-inflammatory, rather than pro-resolution, immune cells (53-55). In addition, PWH with CWP had significantly lower levels of anti-inflammatory, IL-10, compared to HIV negative individuals with chronic low back pain (**Figure 2D**). The plasma concentration of other 6 cytokines were below the reading threshold of the measuring kit and are not reported. Since a prolonged or ongoing presence of pro-inflammatory cytokines can reduce endogenous opioid peptides (25), blood leukocytes were also isolated for measurement of β-END. In leukocytes from PWH with CWP, significantly lower levels of β-END were detected compared to those isolated from HIV-negative individuals with LBP and both no-pain groups (**Figure 2E**). In addition, the mean circulating β-END concentration in plasma of healthy individuals (100 pg/ml) was twice that of pain-free PWH (50 pg/ml) and four-fold higher than that of PWH with CWP (25 pg/ml) (**Figure 2F**). This was not due to diurnal variation in opioid release (56), since blood was collected between 9am-11am to minimize this effect.

**Figure 2.**
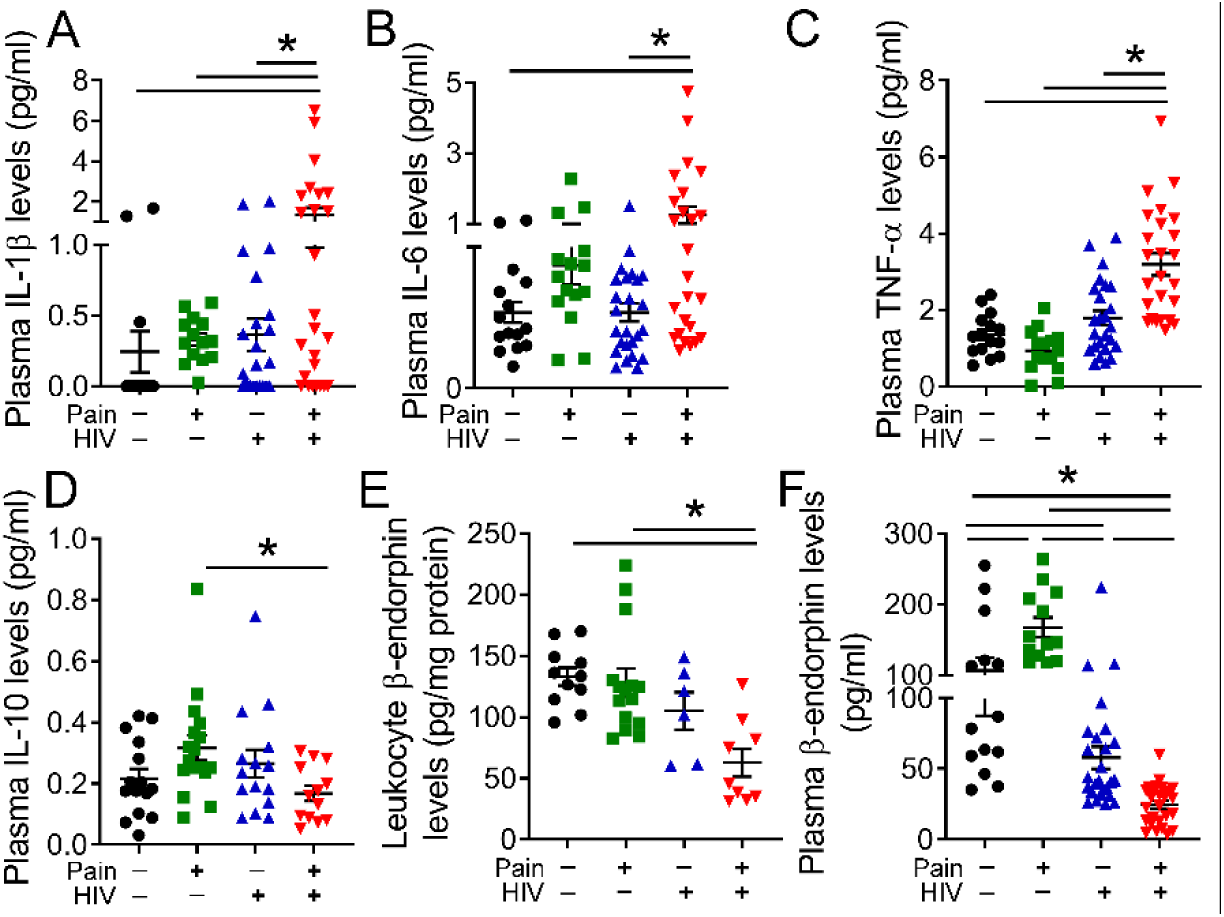
PWH with CWP have increased pro-algetic cytokines and reduced β-END levels. PWH with CWP had significantly higher levels of pro-inflammatory and pro-algetic cytokines in plasma, including IL-1β (n=15-25) (A), IL-6 (n=15-25) (B), and TNF-α (n=15-25) (C). PWH with CWP also had significantly lower levels of anti-inflammatory cytokine, IL-10 (n=13-25) than HIV-negative individuals with LBP (D). Leukocyte levels of β-END were significantly lower in PWH with CWP compared to HIV-negative, pain-free controls and PWH HIV without pain (n=6-15) (E). Circulating β-END levels in plasma were higher in HIV-negative individuals with LBP, but lower in PWH without pain. PWH with CWP had the lowest concentration of plasma β-END (n=13-25) (F). Individual values and means ± SEM. \**P* < 0.05 vs. groups at the end of individual lines; one-way ANOVA followed by Tukey post-hoc testing.

### CWP in PWH is associated with increased ER stress

ER stress is one of the adverse effects triggered by the presence of cell-free heme (32). Given the observed elevation in heme in PWH with CWP, the presence of ER stress was examined in plasma and peripheral leukocytes by measurement of glucose-regulated protein 78 (GRP78/BiP), an important regulator of the unfolded protein response (UPR). GRP78/BiP is the only ER stress marker that is secreted from the cell, allowing it to be measured in both cells and in circulation (32). The results of this experiment demonstrated that PWH with CWP had almost 2-fold higher GRP78/BiP in leukocytes (**Figure 3A**) and 3-fold higher GRP78/BiP in plasma (**Figure 3B**) compared to HIV-negative individuals with pain and both non-pain groups. Since prolonged ER stress causes cell death, plasma activity of lactate dehydrogenase (LDH), a stable enzyme that leaks from cells upon plasma membrane damage (57), was also measured. Consistent with evidence of ER stress, PWH with CWP had significantly elevated plasma LDH activity compared to PWH without pain and HIV-negative controls with or without pain (**Figure 3C**).

**Figure 3.**
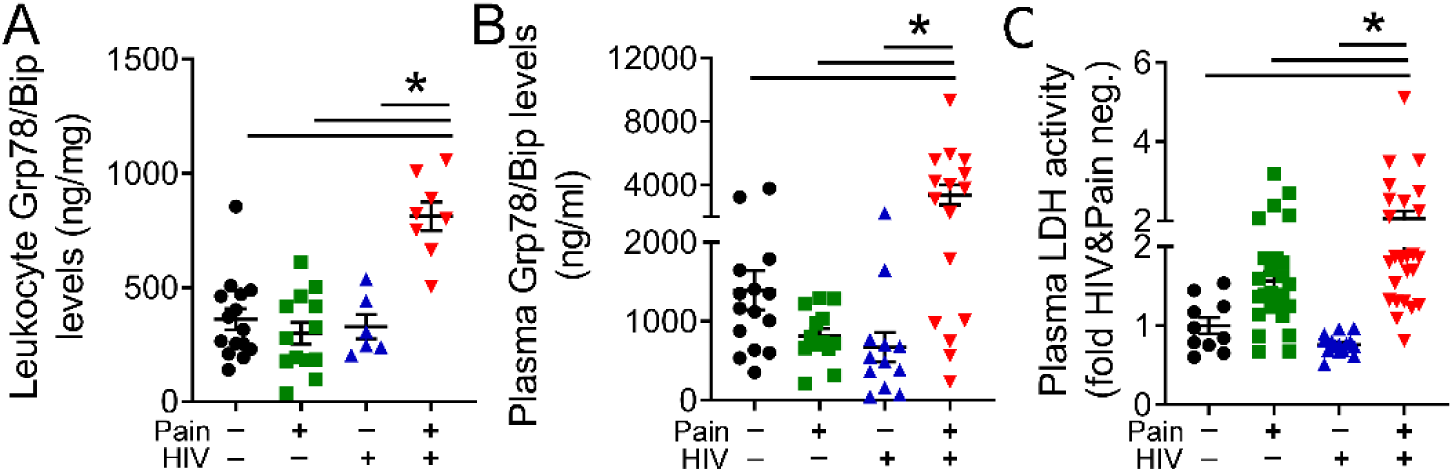
PWH with CWP have increased ER stress. Levels of the ER stress marker, glucose-regulated protein 78 (Grp78/BiP), were significantly higher in leukocytes (n=6-18) (A) and plasma (n=12-21) (B) of PWH with CWP compared to pain-free HIV-negative individuals and pain-free PWH. Lactate dehydrogenase (LDH) activity in the plasma of PWH with CWP was 2-fold higher than in all other groups (n=10-25) (C). Individual values and means ± SEM. \**P* < 0.05 vs. groups at the end of individual lines; one-way ANOVA followed by Tukey post-hoc testing.

### Heme-induced ER stress reduces β-END in immune cells

To establish a direct link between cell-free heme and ER stress with decreased endogenous opioid production, β-END levels/release in murine immune cells (monocyte/macrophage-like J774A.1 cells) was assessed following exposure to hemin (ferric chloride heme) *in vitro*. Compared to DMSO-treated cells, β-END secretion induced by exposure to norepinephrine (NE) was significantly reduced in cells exposed to hemin (**Figure 4A**). This effect was blocked by incubation with sodium 4-phenylbutyrate (4-PBA), a chemical chaperone that reduces ER stress (58). Further, intracellular β-END content was depleted in hemin-exposed cells, but not in cells treated with hemin+4-PBA (**Figure 4B**). Taken together, these results indicate that heme-induced ER stress attenuates β-END levels and release by immune cells.

**Figure 4.**
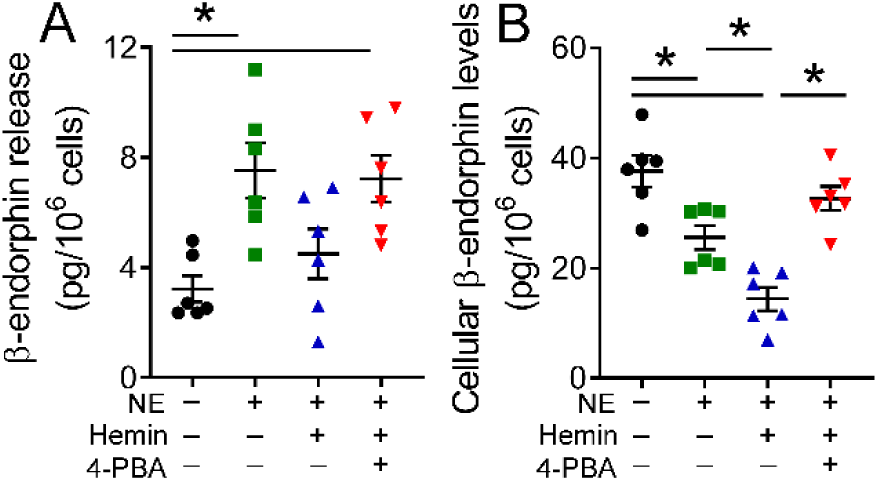
Heme-induced ER stress attenuates β-END levels/release from immune cells. Cultured J774A.1 cells (murine macrophages) were exposed to (hemin, 25 µM, 24 hours) or vehicle (dimethyl sulfoxide, DMSO) in the presence or absence of a chemical chaperone, sodium 4-phenylbutyrate (4-PBA; 10 µM). Norepinephrine (NE; 100nM) was added 5 minutes before the end of incubation to stimulate β-END secretion into the supernatant. NE significantly increased β-END secretion in DMSO-treated, but not in hemin-treated, cells (n=6) (A). 4-PBA restored β-END release in hemin-treated cells. Intracellular β-END levels was depleted in hemin-exposed cells but not in cells treated with hemin + 4-PBA (n=6) (B). Individual values and means ± SEM. \**P* < 0.05 vs. groups at the end of individual lines; one-way ANOVA followed by Tukey post hoc testing.

### Heme scavenging ameliorates disrupted opioid homeostasis and mechanical hyperalgesia in a mouse model of hemolysis

To translate our clinical and *in vitro* findings to an *in vivo* model, phenylhydrazine hydrochloride (PHZ) was used to induce hemolysis in adult male C57BL/6 mice and hindpaw mechanical thresholds were assessed over time. Exposure to PHZ for two consecutive days induced a significant reduction in paw withdrawal thresholds compared to baseline, and this effect was blocked by administration of purified human Hx six hours following the second dose of PHZ (**Figure 5A**). In the same mice, plasma concentrations of cell-free heme were elevated 24 hours following the second dose of PHZ, while the plasma levels of cell-free heme were significantly lower in the mice that received Hx (**Figure 5B**). Using separate cohorts of mice, we found that PHZ-treated mice had elevated levels of Grp78/BiP (**Figure 5C**) and attenuated concentration of β-END in leukocytes (**Figure 5D**) and plasma (**Figure 5E**) relative to saline-treated mice. However, each of these changes were mitigated in mice treated with Hx (**Figures 5C-E**). These results directly link elevated heme with reduced β-END levels and mechanical pain threshold.

**Figure 5.**
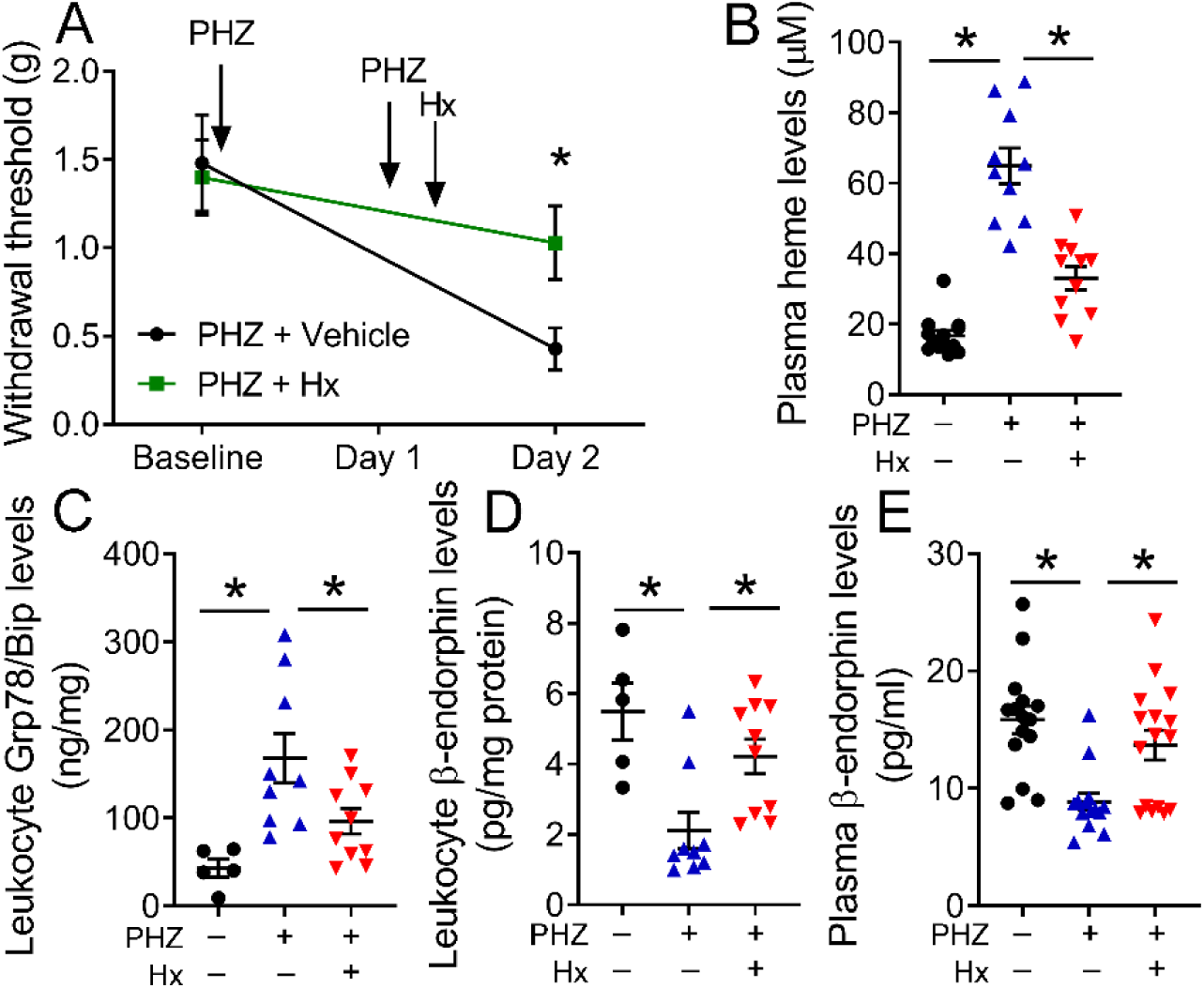
Heme scavenging increases mechanical pain threshold in a rodent model of hemolysis. Phenylhydrazine hydrochloride (PHZ, 50 mg/kg, IP) or saline was administered to adult male C57BL/6 mice on two consecutive days. Six hours following the second injection, subset of mice were given purified human hemopexin (Hx) (4 mg/kg, IP) or saline. Mechanical hypersensitivity was assessed at baseline and on day two by measuring paw withdrawal thresholds to application of von Frey filaments. PHZ-challenged mice exhibited decreased paw withdrawal thresholds compared to mice that received Hx post-PHZ (n=9-10) (A). On day two, mice exposed to PHZ had elevated plasma levels of cell-free heme (n=10-13) (B), higher levels of the ER stress marker, GrpP78/BiP, in peripheral leukocytes (n=5-10) (C) and lower levels of β-END in both leukocytes (n=5-10) (D) and plasma (n=14-16) (E). Hx treatment abrogated the effects of PHZ. Individual values and means ± SEM. \**P* < 0.05 vs. groups at the end of individual lines; unpaired t-tests (A) or one-way ANOVA followed by Tukey post hoc testing (B-E).

### Mice lacking HO-1 exhibit mechanical hyperalgesia and reduced leukocyte β-END

Previous studies have reported that upregulation of HO-1, a heme degrading enzyme, mitigates inflammation- and injury-induced hyperalgesia in mice (43, 44, 59, 60). The underlying mechanisms of this effect are not known. In the present study, we determined whether global knockdown of HO-1 in mice was associated with reduced circulating β-END and a predisposition toward increased nociception. To validate this model, HO-1 protein was measured in mouse liver homogenates, and complete knockdown in HO-1 knockout (HO-1^-/-^) mice was demonstrated (**Figure 6A**). Next, mechanical sensitivity was assessed in HO-1^-/-^ and wild type (WT) mice at baseline and two days after administration of PHZ. Because HO-1^-/-^ mice are unable to metabolize heme and thus highly sensitive to hemolysis with very high mortality, half of the PHZ dose given in previous experiments (i.e., Figure 5) was administered. Interestingly, HO-1^-/-^ mice exhibited significantly lower paw withdrawal thresholds than WT mice at baseline, prior to PHZ treatment (**Figure 6B**). Administration of low-dose PHZ did not further increase mechanical hypersensitivity in HO-1^-/-^ mice, and WT mice did not differ HO-1^-/-^ following PHZ (**Figure 6B**).

**Figure 6.**
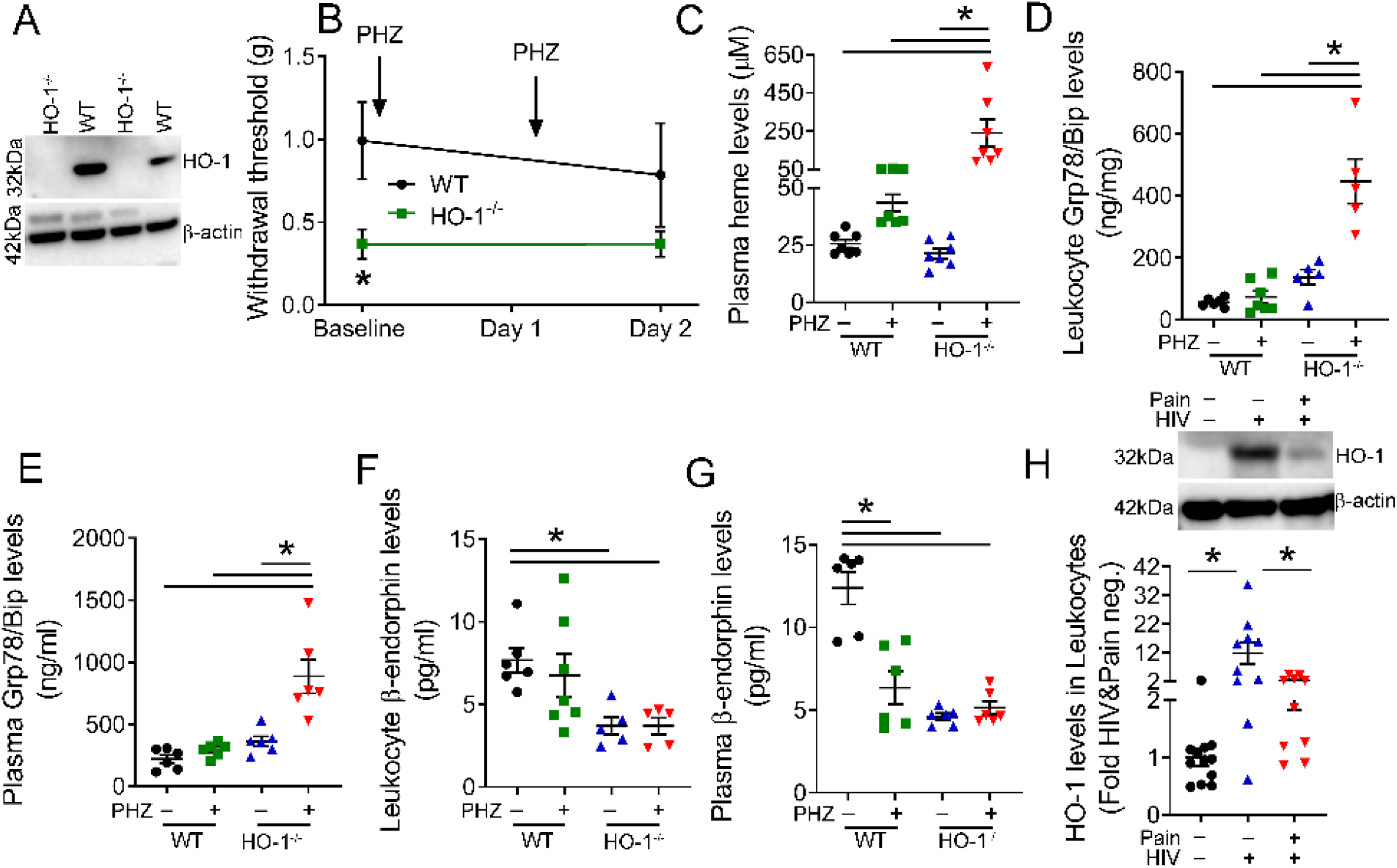
Heme oxygenase-1 (HO-1) knockout mice have low mechanical pain threshold and β-END levels. Immunoblot from liver homogenate validated that HO-1^-/-^mice (adult male) lack HO-1 expression compared to wild type (WT) mice (n=4) (A). Mechanical hypersensitivity was assessed at baseline and after administration of PHZ (25 mg/kg, IP) or saline (vehicle) on two consecutive days by measuring paw withdrawal thresholds to application of von Frey filaments. Paw withdrawal thresholds were lower at baseline in the HO-1^-/-^ mice compared to wild type (WT) mice (n=6) (B). PHZ did not further lower pain threshold in these mice. One day following the 2^nd^ dose of PHZ, HO-1^-/-^ mice had increased plasma levels of cell-free heme (n=7) (C), higher leukocyte (n=5-7) (D) and plasma (n=6) (E) Grp78/BiP levels, and lower leukocyte (n=5-7) (F) and plasma (n=6) (G) β-END levels. HO-1^-/-^ mice thatwere not exposed to PHZ also had low β-END levels (F-G). Immunoblot showed that the expression of HO-1 was significantly lower in the leukocytes of PWH with CWP compared to PWH without pain (n=10-12) (H). Individual values and means ± SEM. \**P* < 0.05 vs. groups at the end of individual lines; unpaired t-tests (B) or one-way ANOVA followed by Tukey post hoc testing (C-H).

Biochemical analysis of plasma and leukocytes from WT and HO-1^-/-^ mice was performed 24 hours following two consecutive daily treatments with PHZ. We observed a significant increase in the plasma concentrations of cell-free heme (**Figure 6C**), and in leukocyte (**Figure 6D**) and plasma (**Figure 6E**) concentrations of Grp78/BiP in HO-1^-/-^ mice, relative to WT mice. Further, the mean plasma concentration of cell-free heme in HO-1^-/-^ mice following 25 mg/kg of PHZ was higher than that of C57BL/6 mice given 50 mg/kg of PHZ (**Figure 5**), likely due to increased sensitivity to hemolysis in the context of HO-1 knockdown. The concentrations of β-END in leukocytes (**Figure 6F**) and plasma (**Figure 6G**) were lower in the HO-1^-/-^ mice irrespective of PHZ treatment, suggesting that HO-1 may regulate β-END independently of heme-induced ER stress. In parallel experiments using leukocytes from PWH, it was demonstrated that the expression of leukocyte HO-1 in PWH with CWP was significantly lower than in PWH without pain (**Figure 6H**), even though those with CWP had significantly higher levels of cell-free heme (**Figure 1A**). Together, these results establish a link between HO-1 expression and β-END levels in leukocytes and suggest that low levels of HO-1 in PWH with CWP may be an additional mechanism independent of heme contributing to hyperalgesia.

## DISCUSSION

This study demonstrates a relationship between self-reported CWP in PWH, with elevated cell-free heme and ER stress markers in plasma, and increased ER stress and diminished levels of β-END in leukocytes. Furthermore, it demonstrates that *in vitro* exposure to heme attenuates β-END content/release in murine monocyte/macrophage-like J774A.1 cells. This effect was reversed by the inhibition of ER stress. Similarly, the induction of hemolysis in C57BL/6 mice increased ER stress and reduced β-END levels in leukocytes, and was accompanied by a decrease in mechanical sensitivity thresholds. This novel mechanism of β-END regulation by cell-free heme-induced ER stress in peripheral immune cells may contribute to hypersensitivity in a subset of PWH.

Plasma levels of cell-free heme are elevated in several hemolytic and chronic inflammatory diseases, such as sickle cell disease (35, 41, 42), lupus arthritis (61), malaria (62), and following treatment with widely used platinum-based anticancer drugs (63). However, the role of heme in HIV associated co-morbidities has not been yet elucidated. There have been few case reports suggesting that hemolysis, heme scavenging, and heme metabolizing mechanisms are impaired in a subset of HIV-positive individuals. For instance, in a recent case report, hemolysis was the main presentation of acute HIV infection in a 22-year-old patient with G6PD-deficiency (64), although the presence or absence of pain in this case was not reported. It is to be noted that the prevalence of G6PD deficiency in HIV-positive individuals is estimated to be 6.8%–13% (65, 66). HIV also confers a 15-40 fold higher risk of acquired thrombotic microangiopathy (45-47), which is an important cause of hemolysis. Further, HIV positive individuals are at risk of hemolysis secondary to the use of ART (48) or other drugs for HIV-associated infections such as the use of amphotericin B and co-trimoxazole for cryptococcal meningitis (49) and use of interferon and ribavirin for hepatitis C (50).

PWH have high reactive species production in peripheral immune cells (67, 68) and elevated lipid peroxidation products in plasma (69). We and others have previously shown that RBCs are especially susceptible to oxidative damage by lipid peroxidation (32, 33, 70), resulting in release of hemoglobin (Hb), metHb, free heme, and free iron (71). In this study we also found that PWH who self-report CWP had elevated levels of carbonyl adducts in RBC membranes. Oxidative stress-induced carbonylation of proteins creates neoantigens against which autoantibodies may develop, as earlier reported in diseases such as COPD (72). Therefore, it is possible that PWH with CWP have these autoantibodies that contribute to hemolysis. These individuals also had increased inflammatory cytokine profiles, suggesting that persistent inflammation may be driving hemolysis. In this regard, environmental and social factors or HIV-associated inflammatory diseases may be responsible for higher oxidative stress seen in HIV-positive individuals with CWP. Moreover, cell-free heme released post-hemolysis is highly inflammatory and may further cause hemolysis, leading to a feed-forward loop of hemolysis-heme-hemolysis (73).

Physiologically, heme concentrations in the blood are maintained at low levels (74) by the high binding affinity of serum heme scavenging proteins, Hx and haptoglobin (75-78). Cell-free Hb binds with circulating haptoglobin, resulting in a heterodimeric complex that is internalized by the transmembrane CD163 receptor on monocytes/macrophages (79). Similarly, cell-free heme binds with Hx and the complex is internalized by the CD91 receptor (80). Once inside the cell, heme and Hb are degraded by HO-1 (81). Immune cells from HO-1-deficient mice are more sensitive to heme-induced oxidative stress (82). A recent case report described impaired CD163-mediated heme scavenging with non-detectable Hx in an HIV-positive individual with severe inflammation (83). In our study, we found that Hx levels in plasma and HO-1 levels in leukocytes were diminished in PWH with CWP compared to the PWH without CWP. These results suggested that the increased hemolysis and diminished heme metabolism are responsible for elevated plasma levels of cell-free heme in PWH with CWP.

We have previously reported that cell-free heme induced cellular toxicity is mediated by ER stress in both animals and humans after bromine gas exposure (32, 33). Accumulation of misfolded proteins in the ER leads to ER stress and activation of the unfolded protein response (UPR) which serves to reduce translation of proteins in order to prevent further accumulation of mutant proteins (84, 85). Recent studies have showed significantly higher levels of misfolded proteins and UPR markers such as Grp78/BiP in people infected with HIV-1 (86-89). We found that the levels of the Grp78/BiP, a master regulator of UPR (90), were higher in HIV-positive individuals with CWP, which corresponded with high levels of cell-free heme in this subset of individuals. However, heme may not be the only inducer of ER stress in PWH. For instance, the HIV-1 transactivator of transcription (Tat) and glycoprotein (gp) 120 proteins have been shown to induce ER stress and cytotoxicity contributing to HIV-associated neuropathogenesis (91, 92). Whether, these HIV proteins increase ER stress independent of cell-free heme or they cause hemolysis and thereby induce heme-dependent ER stress need further investigation.

ER stress has been implicated in inflammatory pain (93) and diabetic peripheral neuropathic pain (94). However, the mechanism by which ER stress modulates pain sensation is unknown. Our *in vitro* analysis showed that heme-induced ER stress attenuated the levels and release of β-END from immune cells. The data also showed that PWH who self-report CWP with high ER stress have 4-fold lower plasma levels of β-END compared to the healthy controls and 2-fold lower β-END than PWH without CWP. We also found that the leukocytes, which are an important source of opioids, have diminished capacity to produce β-END in HIV-positive individuals with CWP. An earlier study compared plasma and brain β-END levels in 48 HIV-positive and 19 health subjects and found that HIV-positive individuals had significant low levels of β-END compared to the controls and that the decrease was not correlated to the CD4^+^T lymphocytes number (95). Similarly, β-END levels in our study did not correlate with CD4^+^ cells as all HIV-positive individuals had low viral load and high CD4^+^ cells. However, the low levels of β-END were inversely proportional to cell-free heme and ER stress.

To determine, whether enhancing heme scavenging by Hx or heme metabolism by HO-1 would increase β-END and attenuate hypersensitivity, we generated an animal model of PHZ-induced hemolysis. Adult, male C57BL/6 mice challenged with the hemolytic agent, PHZ (50 mg/kg body weight), over two consecutive days had a significant increase in plasma concentration of cell-free heme. A similar dosing regimen of PHZ has been shown previously to induce hemolysis and not increase mortality in mice (96). We found that the increase in hemolysis and ER stress correlated with a subsequent decrease in β-END and hindpaw withdrawal threshold in mice. The treatment of the PHZ challenged mice with Hx prevented the decline in β-END levels and the induction of hyperalgesia.

In our HO-1^-/-^ mice, the plasma and leukocyte levels of β-END were significantly lower at baseline compared to the corresponding WT mice which correlated with low paw withdrawal threshold in these mice. At baseline, cell-free heme and ER stress was not elevated in HO-1^-/-^ mice. However, β-END levels were still lower suggesting that HO-1 may directly influence endogenous opioid levels in cells independent of heme. The induction of HO-1 enzyme has been historically associated with reduction in inflammatory pain (44, 97, 98). HO-1 has been shown to potentiate the analgesic effects of morphine (59), probably by increasing the expression of the µ-opioid receptors (99). We found that HIV-positive individuals with CWP had much lower levels of HO-1 compared to HIV-positive individuals without CWP. These low HO-1 levels would not only diminish their ability to metabolize cell-free heme, but also directly reduce β-END levels.

In conclusion, this study has identified a potential novel mechanism of CWP in PWH. Strategies aimed at reducing cell-free heme burden and increasing endogenous opioids may help mitigate CWP in the context of HIV-1 infection. One such strategy was used by Pannell et al. (25), where transfer of macrophages containing higher levels of endogenous opioid was used in mice to ameliorate mechanical hypersensitivity. The analgesic effect was reversed by the perineurally applied opioid receptor antagonist, naloxone methiodide (25). Similarly, peroxisome proliferator-activated receptor-γ (PPAR-γ) agonists, which increase endogenous opioids have been shown to elevate nociceptive threshold in inflammatory pain models (28, 100). Additional therapies to reduce cell-free heme such as Hx, and to some extent albumin and haptoglobin, in addition to pharmacological inhibition of ER stress or induction of HO-1, may also prove beneficial countermeasures against pain in the context of HIV-infection or other hemolytic disorders.

## METHODOLOGY

### Human participants

This study was conducted at the University of Alabama at Birmingham (UAB) and approved by the UAB Institutional Review Board (IRB Protocols 300000860 and 170119003). Participants were categorized as 1) Healthy controls, based on negative HIV status and absence of any pain or chronic disease which may cause hemolysis, 2) HIV-negative with chronic LBP, 3) HIV-positive individuals without any perceived chronic pain, and 4) HIV-positive individuals with self-report of CWP. HIV-positive and -negative participants were recruited from the UAB Center for AIDS Research Network of Integrated Clinical System (CNICS) site. Pain Patient Reported Outcomes (PROs) were part of CNICS as reported earlier (51). The pain PROs consisted of a Brief Chronic Pain Questionnaire about the intensity and duration of the pain. Participants with chronic LBP were recruited via flyers posted at the Pain Treatment Clinic within the UAB Department of Anesthesiology and Perioperative Medicine and the surrounding community. Participants were included if they were active patients at the UAB Pain Treatment Clinic, and reported chronic LBP that had persisted for at least three consecutive months and was present on at least half the days in the past six months (28). Participants were only included if they denied any type of low back surgery or significant trauma/accident within the past year. Low back pain was the primary pain complaint reported for all participants with chronic LBP. Demographic and clinical information was recorded from all participants and blood was drawn. Blood samples were processed, aliquoted, and RBCs and plasma were isolated and stored at -80°C using Freezerworks Sample Inventory Management software (Dataworks Development, Inc, Mountlake Terrace, WA, USA). No samples underwent freeze-thaw cycles prior to use.

### Animals

Adult male C57BL/6 mice (20-25 g) were purchased from Charles River (Wilmington, MA), Heme oxygenase-1 knockout (HO-1^-/-^) mice on a mixed C57BL/6 and FVB background and wildtype (WT) littermates were obtained from Dr. Anupam Agarwal at UAB, details of whichhave been published earlier (101). All mice were housed in conventional polycarbonate cages with woodchip bedding under a 12 hour: 12 hour light/dark cycle with *ad libitum* access to a standard diet and water. Euthanasia protocol based on intraperitoneal injections of ketamine and xylazine was used in the study for mice to minimize pain and distress. All animal care and experimental procedures were approved by the Institutional Animal Care and Use Committee at the University of Alabama in Birmingham (Protocol number: 21416).

### Chemicals

Hemin (ferric chloride heme; product no. H9039), PHZ (for induction of hemolytic anemia; product no. 114715), and norepinephrine (NE, product no. N5785) were obtained from Sigma-Aldrich (St. Louis, MO). Sodium 4-phenylbutyrate (4-PBA; product no. ALX-270-303), a chemical chaperone that reduces ER stress (58), was obtained from Enzo Life Sciences (Farmingdale, NY). Hx (heme scavenger; product no. 16-16-080513) was obtained from Athens Research and Technology (Athens, GA).

### Blood plasma and cell concentration measurements

Heme concentration in plasma samples from humans and mice was measured using the QuantiChrom heme assay kit (product no. DIHM-250; BioAssay Systems, Hayward, CA), according to the manufacturer’s instructions. Plasma concentration of Hx was measured using the human Hx ELISA kit (product no. GWB-4B6D1A; GenWay Biotech, Inc. San Diego, CA. A panel of 10 cytokines [IFN-γ, IL-1β, IL-2, IL-4, IL-6, IL-8, IL-10, IL-12 (p70), IL-13, and TNF-α] were assessed in plasma from human participants using a V-PLEX Proinflammatory Panel 1 (human) cytokine kit (product no. K15049-1D, Meso Scale Diagnostics, LLC, Rockville, Maryland). Cytokine concentrations were measured using a MESO QuickPlex SQ 120 electrochemiluminescence plate reader. Human plasma and mouse J774A.1 cell concentrations of β-END were measured using QuickDetect beta-Endorphin ELISA kits (human: product no. E4458-100; mouse: product no. E4459-100; BioVision, Milpitas, CA). Human plasma and leukocyte and mouse J774A.1 cell concentrations of GRP78/BiP were measured using the GRP78/BiP ELISA kit (product no. ADI-900-214-0001; Enzo Life Sciences). Human LDH activity in plasma was assessed using a LDH cytotoxicity assay kit (product no. 88953, Thermo Scientific, Rockford, IL). All assays were performed following the manufacturers’ protocols.

### RBC fragility assay

Blood was obtained from participants in the presence of an anticoagulant. Plasma was separated and the RBCs were washed with isotonic solution 3 times to remove traces of plasma. RBCs were then re-suspended in normal saline. The RBC suspensions along with 4×4mm glass beads (Pyrex) in DPBS were then rotated 360° for 2 hours at 24 rpm at 37°C. The RBC suspension was then centrifuged at 13,400g for 4 min to separate the intact or damaged cells from the supernatant containing heme/hemoglobin from the lysed cells during this mechanical stress. Free heme/hemoglobin was transferred into a new tube and the absorbance of the supernatant recorded at 540nm as described earlier (52). Subsequently, 100% hemolysis of RBCs was achieved by treating them with 1% Triton x-100 solution. The fractional hemolysis of the sample was then obtained by dividing the optical density of the sample by the optical density of the 100% hemolyzed sample.

### Measurement of protein carbonyl adducts

Protein carbonyl adducts in RBC ghosts were measured as previously described (33). Briefly, RBCs were separated from plasma and hemolyzed with 20 mM hypotonic HEPES buffer. The mixture was centrifuged at 14000xg for 20 min and RBC membrane pellet was dissolved in radio-immuno-precipitation assay (RIPA) buffer (Product number: 89901, Thermo Scientific). Protein was quantified using a BCA kit (product no. 23225, Thermo Scientific, Rockford, IL) and equal amounts of protein (10 µg) were loaded onto each lane of a 4-20% gradient gel. Separated proteins were stained with Amido Black (Sigma-Aldrich, St Louis, MO). The presence of protein carbonyl adducts in RBC ghosts was assessed using an Oxyblot protein oxidation detection kit (product no. S7150; EMD Millipore, Billerica, MA) according to the manufacturer’s protocol. The abundance of protein carbonylation was assessed using densitometry and normalization for protein loading by SDS-PAGE gel quantification.

### Leukocyte isolation from blood

Blood drawn from human participants or mice was mixed with an equal volume of 3% dextran for 30 min to separate leukocyte-rich plasma. Leukocytes containing supernatant was centrifuged (10 min, 1000 rpm, 4°C). The supernatant was discarded, and the remaining RBC and leukocytes were collected. The RBCs were lysed by adding hypotonic solution and leukocytes were re-suspended in 1x PBS/glucose (102) and analyzed immediately.

### PHZ and Hx administration

Mice were challenged with an intraperitoneal injection of PHZ (25 or 50mg/kg BW) or saline (vehicle) on 2 consecutive days. Six hours after the second injection, mice were given a single, intra-peritoneal injection of either saline (vehicle) or Hx in 1x phosphate buffered saline (PBS) (4 µg/kg final concentration). Hx stocks were prepared daily in sterile PBS with injection volumes of 75 µl.

### Cell culture

J774A.1 cells (ATCC), macrophages from adult female BALB/cN mice, were cultured to confluence in DMEM media (product no. 11995-065, Thermo Scientific) and then exposed for 24 hours to hemin (25 µM) or vehicle (dimethyl sulfoxide, DMSO) in the presence or absence of 4-PBA (10 µM). Five min before the end of incubation, NE (100 nM) was added to the cells to induce secretion of β-END into the supernatant.

### Western Blot analysis

Mouse liver and human leukocytes were separately homogenized in RIPA buffer containing protease inhibitors. The samples were sonicated for 3×10 sec on ice in 1.5 ml Eppendorf tubes using an ultrasonic liquid processor and centrifuged at 14,000 g for 20 minutes at 4°C. Protein concentration was measured in cleared supernatants with a BCA assay kit. Equal amounts of protein (25 µg) were loaded in 10% Tris·HCl Criterion precast gels (Product no. 567-1093, Bio-Rad Laboratories, Hercules, CA) and transferred to polyvinylidene difluoride membranes (Product no. 162-0177, Bio-Rad Laboratories) and immuno-stained with an anti-HO-1 antibody (1:1000; Product no. ADI-SPA-896-F, Enzo Life Sciences). Bands were detected by the Protein Detector LumiGLO Western blot kit (Product no. 95059-302, Radnor, PA). Protein loading was normalized by re-probing the membranes with an antibody specific to β-actin.

### Mechanical sensitivity assay

To examine cutaneous mechanical sensitivity, mice were placed on a mesh floor inside a rectangular plexiglass chamber and allowed to acclimate for one hour prior to each test session. Then, mechanical allodynia was measured using the simplified up-down method (103) in which calibrated monofilaments (Stoelting, Dale Wood, IL) were applied to the glabrous skin of each hindpaw to determine a paw withdrawal threshold estimate. Filaments numbered 2 through 9 were used and testing always began with filament 5. If no withdrawal was present for a given stimulus, the next highest value monofilament was applied; if a withdrawal occurred, the following stimulus presentation was the next lowest monofilament value. A total of 5 stimulus presentations occurred. Filament number was converted to force and data are expressed in grams.

### Statistical analyses

Statistical analysis was performed using GraphPad Prism version 7 for Windows (GraphPad Software, San Diego, CA). Data are presented as mean ± SEM. Statistical significance was determined by unpaired t-test for two groups or one-way ANOVA followed by Tukey’s post-hoc test for more than two groups. A value of p < 0.05 was considered significant.

## Authors Contributions

**SA**: Study design, data acquisition, analysis, interpretation, writing of the manuscript, and responsible for quality control; **JD**: Study design, data acquisition, analysis, interpretation, and writing of the manuscript; **IA**: Data acquisition and analysis; **PL, CD, & SM:** Data acquisition; **JSM & BRG**: Study design, recruiting of human participants and participant sample acquisition; **SLH**: Study design, recruiting and acquisition of patient samples, interpretation of the data, and quality control; **SM**: Study design and data interpretation and responsible for quality control.

## Acknowledgements

The authors would like to thank Dr. Gang Liu and Dr. Sami Banerjee for providing J774A.1 cells, and Dr. Anupam Agarwal and Ms. Amie Traylor for providing HO-1^-/-^ and WT mice in support of the study.

